# Efficient clathrin-mediated entry of enteric adenoviruses in human duodenal cells

**DOI:** 10.1101/2023.03.06.531250

**Authors:** Miriam Becker, Noemi Dorma, Dario Valter Conca, Nitesh Mistry, Marta Bally, Niklas Arnberg, Gisa Gerold

## Abstract

Enteric adenovirus types F40 and 41 (EAdVs) are a leading cause of diarrhea and diarrhea-associated death in young children and have recently been proposed to cause acute hepatitis in children. Unlike other adenoviruses, EAdVs exhibit hitherto a strict tropism for gastrointestinal tissues with, to date, unknown infection mechanism and target cells. In this study, we turn to potentially limiting host factors by comparison of EAdV entry in cell lines with respiratory and intestinal origin by cellular perturbation, virus particle tracking and transmission electron microscopy. Our analyses highlight kinetic advantages in duodenal HuTu80 cell infection and reveal a larger fraction of mobile particles, faster virus uptake and infectious particle entry in intestinal cells. Moreover, EAdVs display a dependence on clathrin- and dynamin-dependent pathways in intestinal cells. Detailed knowledge of virus entry routes and host factor requirements is essential to understand pathogenesis and develop new countermeasures. Hence, this study provides novel insights into the entry mechanisms of a medically important virus with emerging tropism in a physiologically relevant cell line.

**Author Summary:** Enteric adenoviruses have historically been difficult to grow in cell culture, which resulted in lack of knowledge of host factors and pathways required for infection of these medically relevant viruses. Previous studies in non-intestinal cell lines showed slow infection kinetics and generated comparatively low virus yields compared to other adenovirus types. We suggest duodenum derived HuTu80 cells as a superior cell line for studies to complement efforts using complex intestinal tissue models. We show that viral host cell factors required for virus entry differ between cell lines from distinct origins and demonstrate the importance of clathrin-mediated endocytosis.

## Introduction

Human enteric adenovirus type 40 and 41 (EAdV) are the only human members of species F of the growing *Adenoviridae* family, which currently has over 110 members divided in species A-G (1,2). While other adenovirus types cause disease in eyes, airways, liver, adenoids and/or urinary tract, EAdVs have a pronounced tropism for the gastrointestinal tract (3) and are one of the leading causes for acute virus gastroenteritis and associated death in children below 5 years of age (4,5). In 2022, more than 1,000 cases of acute hepatitis of unknown cause occurred in children worldwide, including 22 deaths (6). In UK, adenoviruses were detected in 122 of 183 cases (69%) and, with human adenovirus type 41 (AdV41) being the most common (92.3%) adenovirus type detected in blood (7), suggesting a potential novel role for these viruses.

Despite being a common cause of disease in humans, the interplay of host factors and virus entry requirements for EAdVs is largely unknown. The main reason for this is that EAdV have been difficult to isolate and amplify to high viral titer in cell culture (8,9). The icosahedral capsid of adenoviruses consists of three capsid proteins, where the major capsid protein, hexon, is the main structural component. Pentamers of viral penton proteins are located at the tips of the icosahedral vertices and from their centers, trimeric fiber proteins are protruding (10). In contrast to other AdVs, EAdVs carry two distinct fiber proteins of different length (11,12), which mediate interaction with surface attachment factors and binding receptors, heparan sulfate proteoglycans (HSPG) and coxsackie- and adenovirus receptor (CAR), respectively (13,14). All other AdV types interact with cell-surface integrins via an RGD-motif in the penton protein, which aids in endocytic uptake and uncoating of the virus (15,16). EAdVs, however, lack such conserved RGD-motifs and therefore interact with laminin-binding integrins instead (17). The overall EAdV structure is largely resistant to pH changes that resemble passage through the stomach, which may contribute to the gastrointestinal tropism (18,19). Due to their fastidious nature, most studies have been conducted in A549 and HEK293 cells where EAdV entry was slow, occurred in a clathrin-independent manner, and with inefficient endosomal escape (20,21). Previous attempts to study EAdV infection in human ileal organoids succeeded in virus amplification but failed to identify EAdV target cells (22). This highlights the need to identify the molecular determinants responsible for the specific EAdV tissue tropism by exploring novel infection models.

In this study, we compared the EAdV infection kinetics and cell entry pathway in duodenal HuTu80 cell lines to lung-derived A549 cell lines to identify possible cell type specific advantages in cells derived from the gastrointestinal tract. We found that infectious internalization was faster and clathrin-dependent in HuTu80 cells highlighting the need for tissue specific virus infection studies. Our results suggest that the clinically relevant but understudied species F adenovirus carry traits not seen in other adenoviruses, which enable efficient entry in cells of the gastrointestinal tract.

## Results

### Faster uptake and infection kinetics of EAdVs in duodenal HuTu80 cells

With the aim to explore the host cell requirements of enteric adenoviruses (EAdVs) in duodenal HuTu80 and lung-derived A549 cells, we confirmed susceptibility of both cell lines to AdV40, AdV41, and AdV5 (S1 Fig.A-C). To study AdV entry in more detail, we established an uptake assay based on dual-labeling of virus particles (Fig. 1A). In brief, AF488 fluorescently labeled AdVs were bound to cells at 4°C followed by a shift to 37°C to allow internalization. To distinguish intracellular and extracellular virus particles, cells were shifted back to 4°C to inhibit further internalization, and extracellular particles were visualized with an anti-AF488 primary antibody and AF568 secondary antibody without fixation. Thereafter live cells were analyzed by flow cytometry (Fig. 1C-E) for total virus signal (AF488) and extracellular fraction (AF568) or fixed and stained for microscopy analysis (Fig. 1G and Fig. 1H). To validate the assay specificity, fluorescently labeled AdV40 was bound to and incubated with cells at 4°C or allowed to directly internalize for 4 h at 37°C before staining with subsequent fixation (Fig. 1B). Analysis by confocal microscopy revealed that extracellular particles were double stained (Fig. 1B top row), whereas internalized particles were protected from anti-AF488 staining (Fig. 1B bottom row). EAdV were described to internalize slowly in A549 and HEK293 cells as compared to other AdV types (20,21). We confirmed by flow cytometry that uptake of EAdV into A549 cells was slow and reached 90-100% after 2-3 h, whereas 75% of AdV5 particles were already taken up after 30 min (Fig. 1C-E). We observed that both AdV40 and AdV41 entered HuTu80 about twice as fast than into A549 cells. In contrast, AdV5 entered into HuTu80 and A549 cells equally fast (Fig. 1E). This suggested a cell line specific advantage for EAdVs in HuTu80 cells with regard to virion internalization.

**Figure 1:**
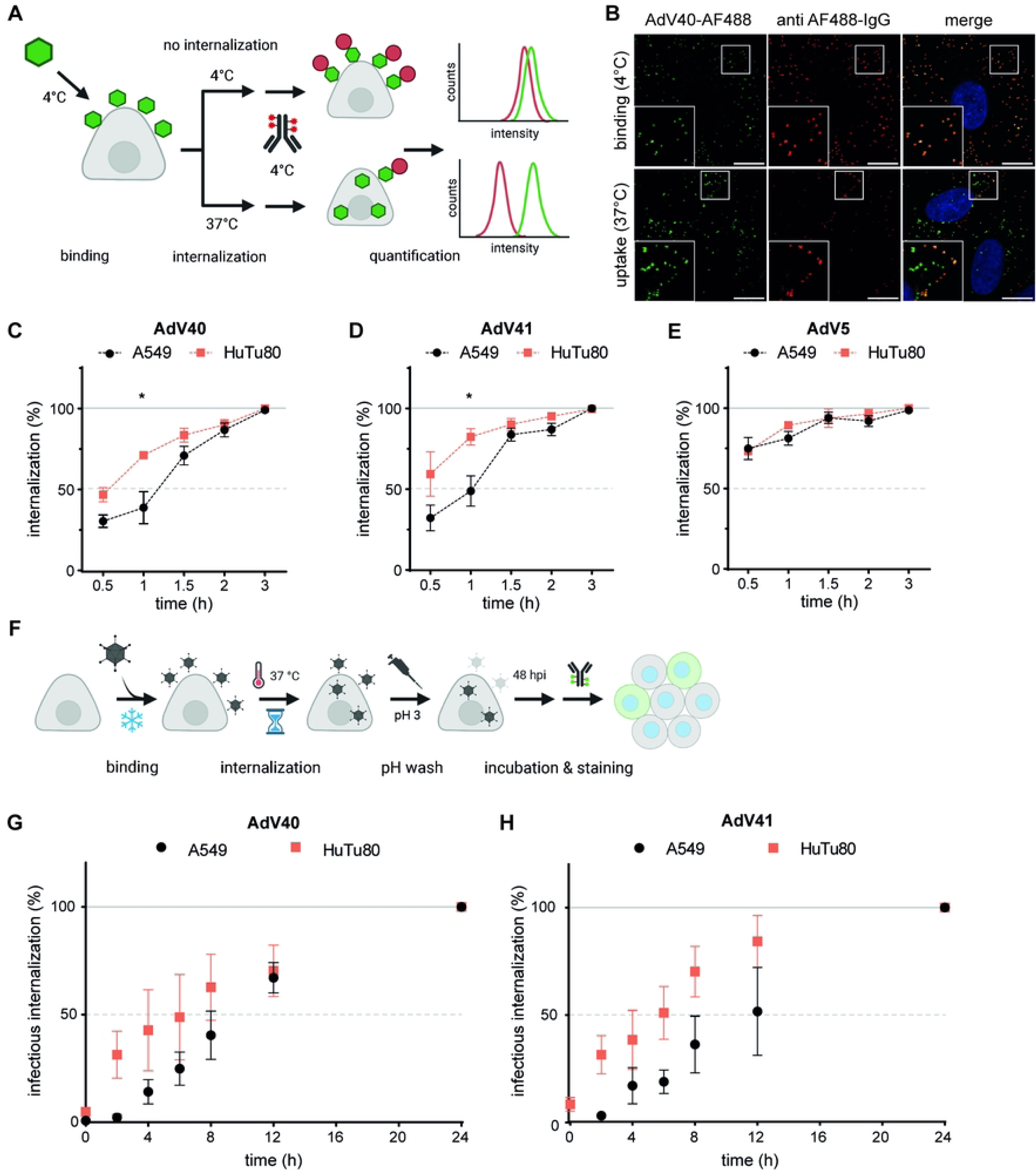
Uptake and infection kinetics of EAdV is faster in duodenal HuTu-80 cells. A) Scheme of the dual label uptake assay to distinguish internalized from extracellular AdVs. AF488-AdVs were bound to target cells in suspension followed by internalization. Extracellular virus accessible to anti-AF488 antibody staining showed a dual labeling measurable by flow cytometry. B) Validation of uptake of AF488-labeled AdV40 bound to A549. Dual staining depended on extracellular localization as in the top panel with the binding only control. Scale bar: 10 µm. C-E) Uptake assay of AdV40 (C), AdV41 (D), and AdV5 (E) in A549 (black) and HuTu80 (red) cells assessed through shifts in mean fluorescence intensities by flow cytometry. F) Scheme of an infectious internalization assay to determine kinetics of infectious uptake of AdVs. G/H) AdVs were bound to and internalized into A549 or HuTu80 cells. Extracellular EAdV particles washed at low pH, whereby only internalized particles led to infection. Error bars show standard error of the mean. Statistical significance by unpaired students t-test: *: p<0.05.

To investigate the relevance of this advantage for productive virus uptake, we next developed an infectious uptake assay, where EAdVs were bound at 4°C, then allowed to enter cells at 37°C for different durations. After the respective internalization time, we rendered extracellular particles non-infectious by low pH wash (S2 Fig.) and quantified successful viral hexon production as a correlate of productive entry 48 h post infection. Both AdV40 and AdV41 reached higher infection levels in HuTu80 cells as compared to A549 cells, especially after washes at early time points between 2 h and 6 h of internalization (Fig. 1G/H). Thus, infectious EAdV internalization was faster in HuTu80 cells as compared to A549 cells. These results suggest that duodenal cells are a more suitable cell model to study EAdV entry than previously used cell lines.

### EAdVs display heterogeneous motion and increased free diffusion with time and in HuTu80

After establishing that EAdV entry is significantly faster in HuTu80 cells than in A549, we investigated if virus particle motion after attachment to the cell surface also differed between cell lines. To this end we tracked single particles of fluorescently labeled AdV by highly inclined and laminated optical sheet (HILO) microscopy (23). We incubated A549 and HuTu80 cells with AF488-AdV5/40/41 for 5 min at 37°C and observed virus motion on cells at 5 min and 30 min after removal of virus inoculum. We acquired videos at 10 frames/s for two min each. The tracks were analyzed using a divide-and-conquer moment scaling spectrum (DC-MSS) algorithm (24,25), which allowed us to subdivide each track into segments displaying distinct motion patterns. Four motion types were considered in the analysis: immobile particles, confined diffusion, free diffusion, and directed motion. Representative examples of the acquired tracks are shown in Fig. 2A. We first analyzed the diffusion coefficient for each motion type. Surprisingly, no clear differences between AdV types or cell lines were observed (S3 Fig.). Both free and confined diffusion had a mean diffusion coefficient between 0.008 and 0.02 µm^2^/s, while directed motion exhibits slightly higher values between 0.015 and 0.035 µm^2^/s. These values were comparable to values reported previously for AdV2 internalization (16). In all cases and for all motion types, we observed large variations in the diffusion coefficient values, with standard deviations >0.02 µm/s and single recorded values ranging between 5×10^-5^ to 0.05 µm^2^/s. This heterogeneity in the diffusion coefficient value can be attributed to the different possible interactions between cell components and the virus, which result in various levels of mobility.

**Figure 2:**
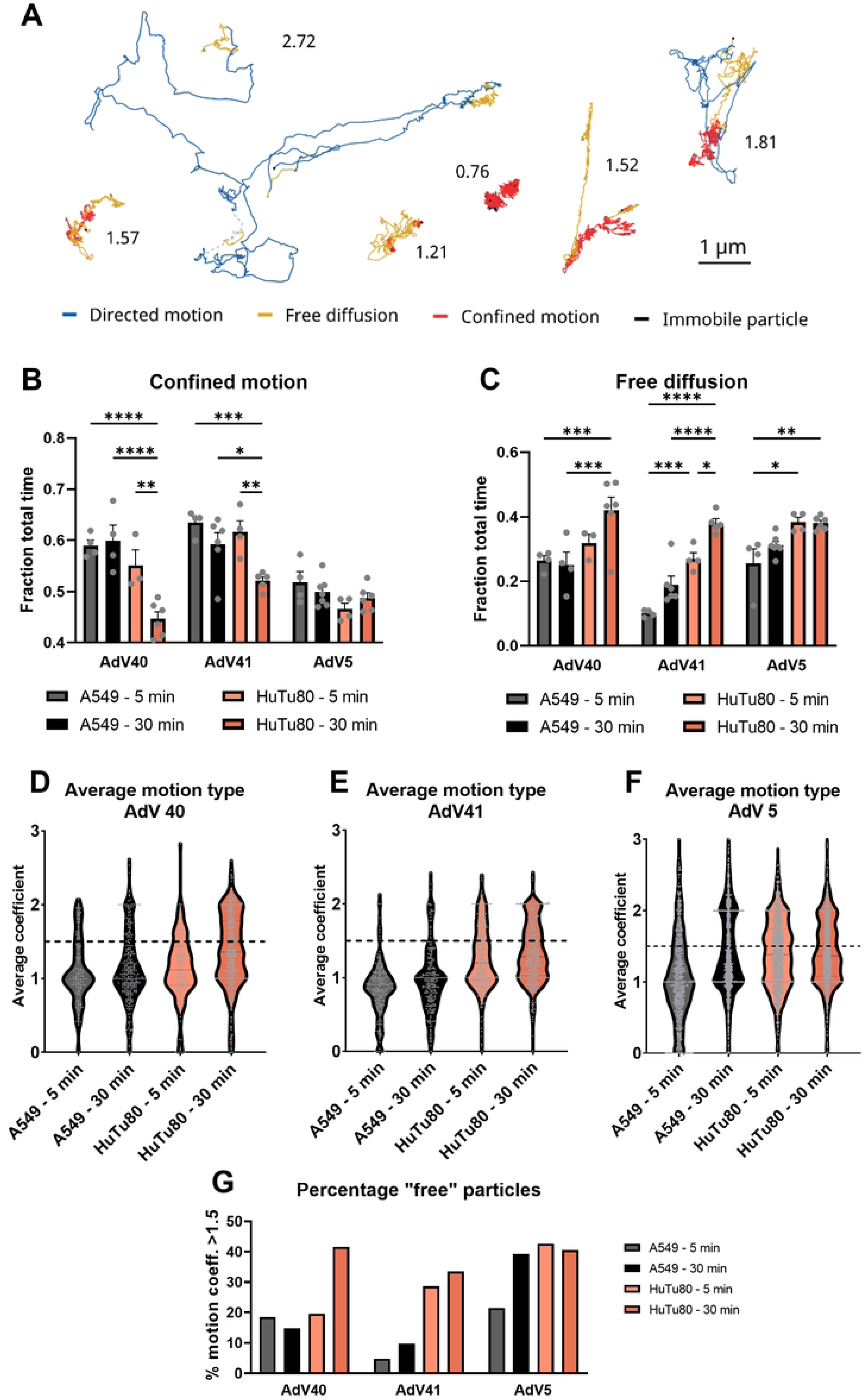
EAdVs display increased free diffusion with time and on HuTu80 cells. A) Representative tracks produced by AdV particles moving on live cells. The colors show the results of the track segmentation algorithm. The number next to each track is its average motion coefficient ranging between 0, for a fully immobile track, and 3, for directed motion. B-C) Fraction of the total track time spent by AdV particles in confined (B) and free (C) diffusion, after 5 and 30 min from the removal of the inoculum and on A549 and HuTu80. Grey dots indicate the value for each recorded video. Error bars indicate the standard error of the mean. Statistical significance determined using two-way ANOVA test: *: p<0.05, **: p<0.01, ***: p<0.001, ****: p<0.0001. D-F) Truncated violin plot showing the distribution of full tracks according to their average motion coefficient on A549 and HuTu80 cells after 5 and 30 min from the removal of the inoculum for AdV40 (D), AdV41 (E) and AdV5 (F). Single tracks are shown as grey dots. G) Percentage of the tracks in (D-F) with an average motion coefficient higher than 1.5, i.e. displaying a motion dominated by free and directed diffusion.

We then focused on the relevance of each motion type, measured as the fraction of time spent by the particles in each of those. We observed in all cases a prevalence of confined motion (from 40% to 68% of the total time) followed by free diffusion and immobile particles, while directed motion only relegated to a few percent of the total time tracked (S4 Fig. A). Noteworthy, the analysis of the fraction of total time likely underestimates the importance of fast but short-lived motion segments. In these segments, AdV particles might cover large distances making them significant in the overall dynamic of the interaction with the host cell. To further explore this possibility, we considered the distance travelled by the particle in each segment as the distance between the two furthest points in each track segment (as visualized in S4 Fig. D/E). In this case, free diffusion becomes the most prevalent motion type, and the importance of directed motion becomes evident, accounting for as much as 20% of the distance travelled by AdV5 particles on A549 cells (S4 Fig. F). We then compared the variations in the frequency of each motion type between cell lines and time points for all three viruses. The behavior of AdV5 shows little dependence on time and cell type, with a small increase of free motion in HuTu80 cells as compared to A549 and a respective reduction of immobile particles (Fig. 2B and S4 Fig. B). This time-independence supports the internalization data in Fig. 1 indicating a fast kinetics for AdV5 internalization regardless of the cell type. Particularly noteworthy is the behavior of AdV40 and AdV41. As shown in Fig. 2B-C, when comparing motion on A549 and HuTu80, we observe a significant increase in free diffusion (from a fraction of 0.25 to 0.42 and from 0.19 to 0.38 after 30 minutes for AdV40 and AdV41, respectively) and reduction in confined motion (from 0.60 to 0.45 and from 0.59 to 0.52 for AdV40 and AdV41). For AdV41, we also observe a clear reduction of immobile particles, which is not present for AdV40 (S4 Fig. B). This indicates a much more dynamic behavior of EAdV particles on intestinal cells. In addition, we observed a significant increase in mobility with time for both AdV40 and AdV41 on HuTu80 cells. The free motion fraction increased by around 10% of the total time for both EAdVs, reaching similar levels measured for AdV5 (Fig. 2B), while the confined fraction was reduced simultaneously (Fig. 2C). We did not observe a significant variation for directed motion (S2 Fig. C), most probably due to the reduce sample size. In sum, we show that EAdVs show an overall increased movement dynamics during the attachment and entry process in intestinal cells.

Combining all segments of a motion type from all tracks, however, does not consider the possible heterogeneity in the diffusion behavior and mobility between single particles. To investigate this, we developed a parameter that describes the overall dominant motion type for each track. We scored every motion type with an integer number between 0 and 3 in increasing order of mobility and defined the “average motion coefficient” (AMC) as the time average of the motion types displayed by each track. In this way, tracks receive a fractional number between 0 and 3, with values close to 1 indicating a predominantly immobile/confined motion and around 2 for free diffusion and directed motion. Examples of AMC scores are shown in Fig. 2A, while the distribution of the AMC for all tracks is shown in Fig. 2D for AdV40 and Fig. 2E for AdV41. In all conditions, a population is present around a value of 1, comprising the particles that display almost exclusively confined motion. This is the only major peak present for A549 cells, with only a small tail reaching AMC values close to 2. Particles on HuTu80 cells instead are characterized by a more heterogeneous track distribution, which is again dependent on the observation time. After 5 min incubation, only one major peak is present around 1, however, the distribution appears significantly broader towards higher scores than in the A549 case. At 30 min a second population appears centered around 2 for both EAdVs. This indicates that rather than a homogeneous increase of the free diffusion mode shared by most particles, the larger free-diffusing fraction is due to the appearance of a population of mostly freely diffusing particles, while others remain immobile or restricted in their motion. For AdV5, we observe a broad distribution extending in into large AMC values in all conditions and the presence of the second peak around an AMC value of 2 in all cases except for A549 cells after 5 min (Fig. 2F). Finally, we estimated the size of the two populations by a simple thresholding at an AMC score of 1.5. After 30 min the number of tracks undergoing mostly free diffusion is 2.8 times higher for HuTu80 than for A549 in the case of AdV40 and 3.4 times higher for AdV41. No clear increase is observed for AdV5 when comparing the two cell lines (Fig. 2G). At the early time point (5 min), the difference between the two cell lines is even larger (6-fold) for EAdV41, while AdV40 shows little difference between the two cell lines and AdV5 a 2-fold increase. Taken together, this indicates that the faster internalization observed for EAdV on HuTu80 cells correlates with an increase in overall mobility. This in turn is driven by the emergence of a highly mobile population, which increases with time.

### Kinetic advantage is reflected in intracellular localization of EAdVs

The observation that EAdVs entered HuTu80 cells rapidly, prompted us to investigate EAdV localization in HuTu80 (Fig. 3B) versus A549 (Fig. 3A) after 12.5 min, 2 h and 6 h of internalization by transmission electron microscopy. We consistently observed virus particles binding to flat membrane regions or at the tip of cellular protrusions (Fig. 3A/B “binding” and Fig. 3C blue) at all time-points and for both cell lines. Moreover, virus particles associated with curved membranes with and without clathrin coat were observed in all experimental conditions (Fig. 3A/B “endocytic pit” and Fig. 3C green); virus particles were also found in intracellular vesicles or endosomes (Fig. 3A/B “endosome” and Fig. 3C red). In rare cases virus particles appeared without endosomal membrane coat in the cytosol indicating successful endosomal escape, however these events were not observed in A549 cells at 12.5 min of internalization (Fig. 3A/B “cytosolic” and Fig. 3C yellow). Quantification of these four virus localizations (plasma membrane, endocytic pit, endosome, and cytosol) confirmed that at these early time points a greater fraction of viruses localized to intracellular compartments of HuTu80 as compared to A549 cells. This was most obvious at 2 h internalization, where only 15%-25% of AdV40 and AdV41 were found intracellularly in A549 cells, but 40%-60% in HuTu80 cells (Fig. 3D vs. Fig. 3E at 2 h). These results are well in line with our observations in flow cytometry and infection assays (Fig. 1). In summary, our data suggest that duodenal HuTu80 cells provide a cell type specific advantage for EAdV infectious entry and may serve as an improved cell culture model.

**Figure 3:**
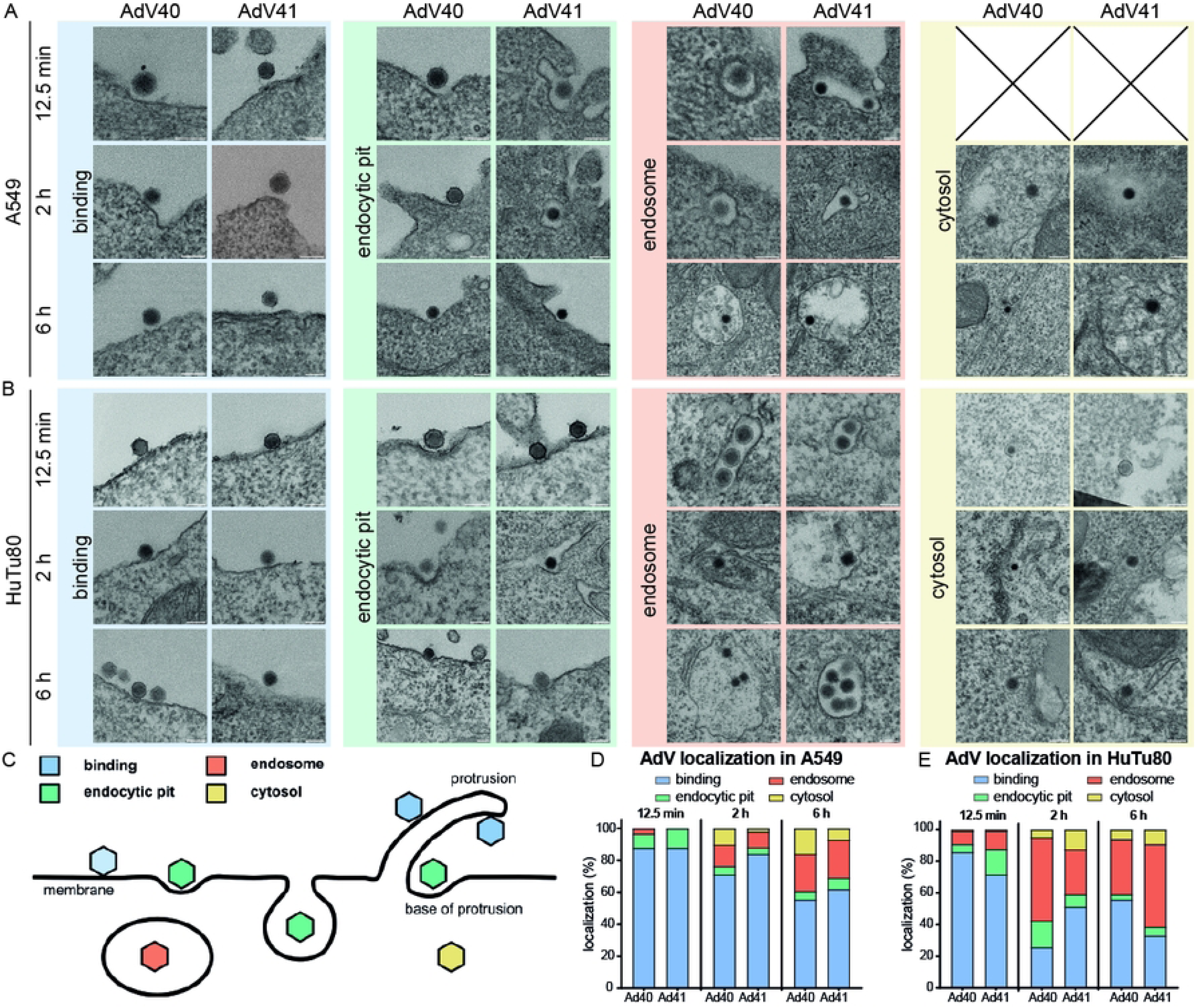
EAdV uptake observed in transmission electron microscopy mirrors kinetic advantage. A) Panel of exemplary cropped TEM images from A549 cells incubated with AdV40 (left) or AdV41 (right) and classification of virus localization into categories: bound (blue), endocytic pit (green), internalized (red), and cytosolic (yellow); B) Panel of exemplary TEM images as in A) from HuTu80 cells. Scale bars: 100 nm. C) Scheme for scoring of localization of AdV particles on cells in the EM experiments; D/E) Quantification of uptake of EAdVs into A549 (D) and HuTu80 (E) cells. At least n=50 virus particles were classified per time point.

### EAdVs enter cells in a clathrin- and actin-dependent manner

Enteric adenovirus entry was previous studied in A549 and HEK293 cells and has been described to be clathrin-independent in HEK293 (20,21). The faster EAdV-specific uptake kinetics in duodenal HuTu80 cells prompted us to study their cell entry pathways using the flow cytometry-based uptake assay (Fig. 1A) in presence of small molecule inhibitors of endocytic pathways. Cells were detached with PBS/EDTA and suspended in growth medium with small molecule inhibitors for 30 min pretreatment. We next added virus particles to cells for 1 h at 4°C in presence of the inhibitors. Thereafter, the virus inoculum was removed and cells were resuspended in medium with inhibitor and shifted to 37°C for internalization for 4 h. We used an anti-AF488 antibody staining on live cells at 4°C as above, to detect remaining extracellular virions.

Clathrin-mediated endocytosis (CME) is a well characterized endocytic pathway (26) and used for cell entry by several AdVs (27). Pitstop2 blocks attachment of clathrin triskelions to the forming endocytic pit (28) and thereby inhibits CME. In our virus entry assay, pitstop2 reduced uptake of EAdVs dose-dependently to a maximum of 50% internalization in HuTu80 cells but interestingly did not affect uptake into A549 cells (Fig. 4A/B). These findings suggested a possible difference between entry pathways in A549 and HuTu80 and prompted us to explore the involvement of CME further. To that end we tested the involvement of dynamins, which are GTPases acting as important scission factors during endocytic vesicle formation in CME, but also during caveolae-dependent endocytosis and phagocytosis (29). We assessed the involvement of dynamins in EAdV endocytosis by inhibition of its GTPase function by dynasore (30). Dynasore inhibited EAdV uptake dose-dependently in both cell lines. Specifically, dynasore reduced internalization rates to below 50% residual uptake for all viruses at the highest inhibitor concentration (Fig. 4C/D). As dynasore treatment affects several pathways, the larger reduction in internalization observed as compared to pitstop2 indicates that several endocytic pathways are active in parallel during uptake.

**Figure 4:**
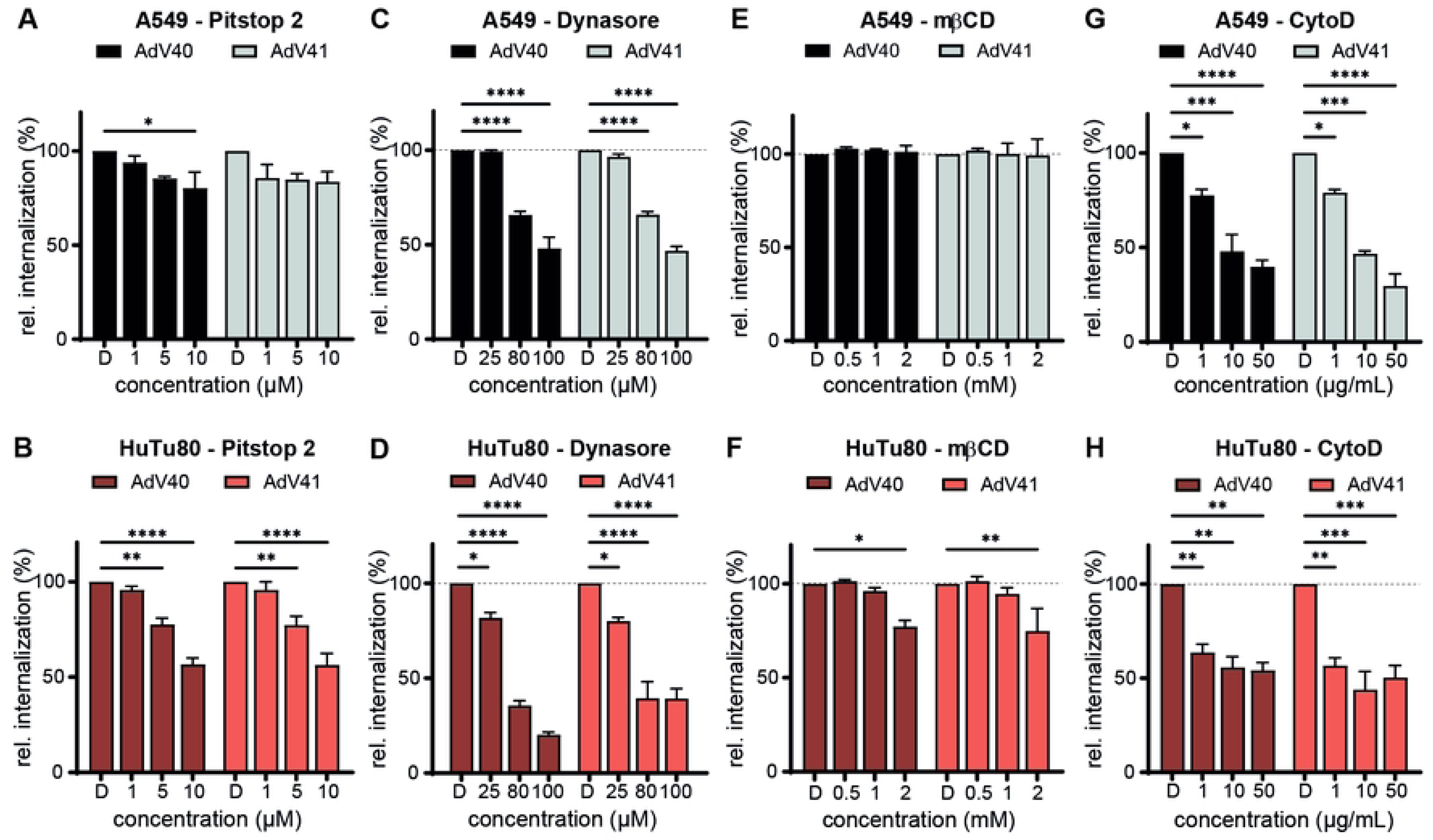
EAdV uptake depends on clathrin, dynamins and actin. Relative EAdV uptake in A549 (black) and HuTu80 (red) cells in the presence of clathrin-inhibitor pitstop-2 (A/B), dynamin-inhibitor dynasore (C/D), cholesterol-depleting agent methyl-beta-cyclodextrin (E/F), with actin-perturbating cytochalasin D (G/H) at indicated concentrations, compared to vehicle control (DMSO). Error bars indicate the standard error of the mean. Statistical significance determined using two-way ANOVA test: *: p<0.05, **: p<0.01, ***: p<0.001, ****: p<0.0001.

Next, we assessed whether EAdV entry required membrane cholesterol in addition to clathrin and dynamin. Cholesterol is an important membrane organizer and its depletion can affect membrane dynamics and domain formation (31,32). We used methyl-β-cyclodextrin (mβCD) to sequester cholesterol from membranes (33,34), which only mildly affected EAdVs uptake (Fig. 4E/F). This suggests that cholesterol-rich domains or cholesterol-dependent membrane dynamics are not involved in EAdV uptake.

Finally, we probed for a role of actin in EAdV entry. Actin dynamics affect, among other processes, the maintenance of membrane rigidity, cell movement and intracellular transport (35). We utilized cytochalasin D (cytoD) to perturb the dynamics of the actin cytoskeleton by blocking actin filament elongation (36). Perturbation of dynamic actin remodeling by cytoD reduced EAdV uptake dose-dependently to maximally 50% residual particle uptake in both cell lines (Fig. 4G/H). This finding is expected, as actin regulates several cellular processes and thus can affect virus uptake at several stages.

In summary, our data suggest that EAdV entry is clathrin-independent in A549 cells, but clathrin and dynamin dependent in HuTu80, pointing towards CME as the major uptake route in duodenal cells. Therefore, we decided to corroborate our findings on particle uptake by testing if infectious EAdV entry was clathrin- and actin-dependent.

Consequently, we established an infection setup optimized for cell survival and stable EAdV hexon readout. Due to the slow infection life cycle of EAdV, stable hexon signal was not detectable before 44-48 h post infection. We treated HuTu80 and A549 cells with pitstop2 and cytoD for 30 min before addition of AdV40 or AdV41. The cells were then incubated for 2 h at 37°C until the inoculum was exchanged for medium containing inhibitor. The inhibitor was removed at 24 h post infection to increase cell viability within the assay and cells were fixed and stained for viral hexon and nuclei at 48 h post infection. Interestingly, pitstop2 reduced EAdV infection of HuTu80 and A549 cells only at the highest inhibitor concentration, but not in a dose-dependent manner (Fig. 5B/C). At 5 µM pitstop2, infection of both cell lines with AdV40 and AdV41 was reduced by more than 50% compared to the DMSO treated control. In contrast, cytoD treatment efficiently and dose-dependently reduced infection to about 25% residual infection at the highest drug concentration for HuTu80 and 50% residual infection for A549 (Fig. 5D/E). From these data we conclude that not only overall particle uptake but also infectious EAdV uptake in HuTu80 was clathrin- and actin-dependent.

**Figure 5:**
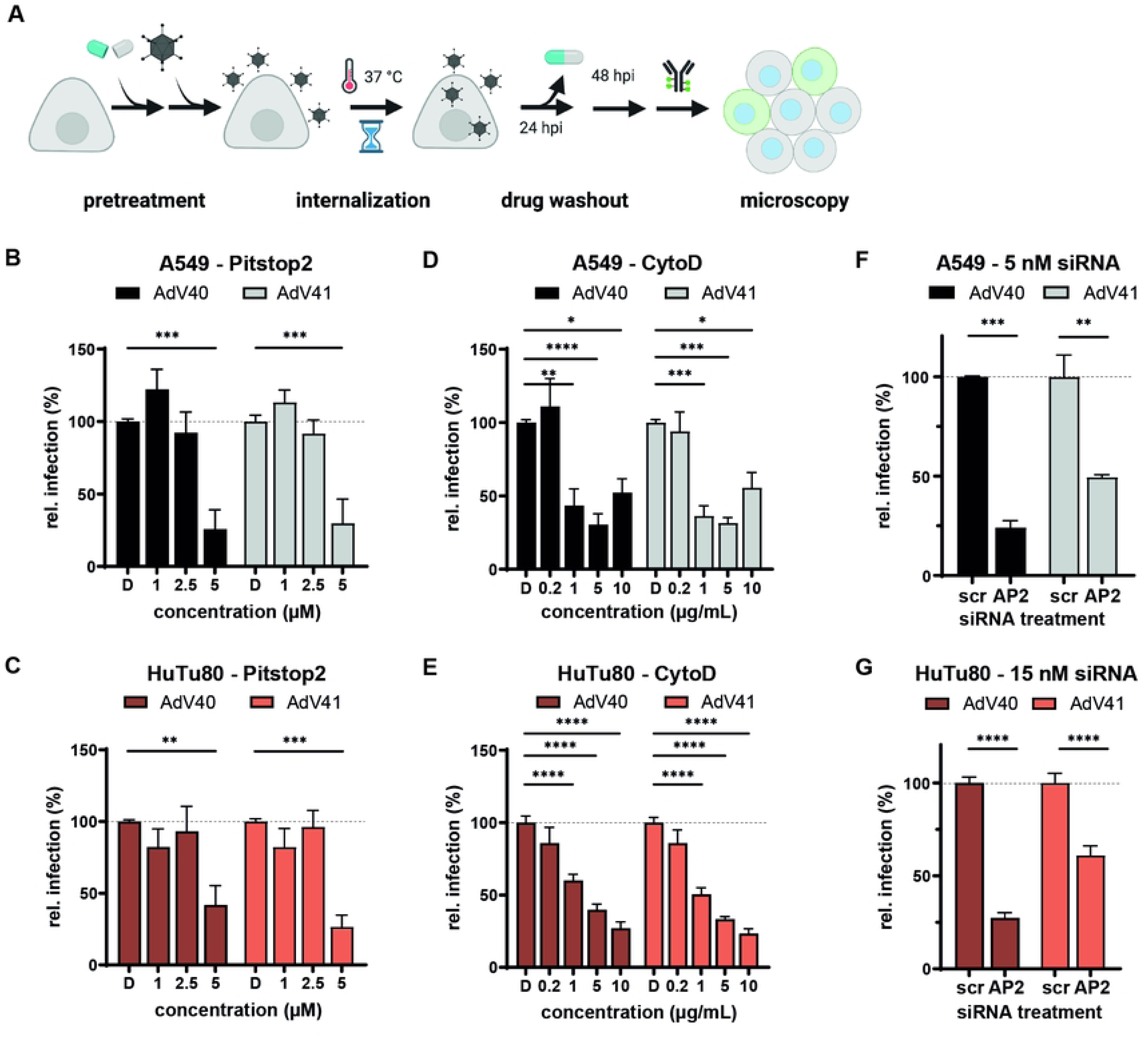
Infectious internalization of EAdV depends on clathrin in HuTu80 cells. A) Scheme of the infectious internalization assay in presence of inhibitors. B-E) Infectious internalization in the presence of pitstop2 (B/C), cytochalasin D (D/E) in A549 (black) and HuTu80 (red) cells. Infectious internalization after treatment of cells with siRNA against clathrin adapter AP2 or non-targeting siRNA scrambled control. Error bars indicate the standard error of the mean. Statistical significance determined using two-way ANOVA test: *: p<0.05, **: p<0.01, ***: p<0.001, ****: p<0.0001.

To verify our findings omitting pitstop2, we used RNA interference against AP2, a clathrin adapter at the plasma membrane (37), followed by infection with EAdV. To generate an efficient knock-down level, we depleted AP2 in HuTu80 and A549 cells by double transfection with 48 h in between with siRNA against AP2 or with a scrambled control (38). We seeded cells 24 h after the second siRNA treatment, infected them 24 h later as stated above and fixed and stained for hexon protein and nuclei at 48 h post infection. We harvested additional cells at the time of infection to determine knock down efficiencies on protein level. Knock down of AP2 reduced infection of HuTu80 by 75% for AdV40 and 40% for AdV41 and infection of A549 by 75% for AdV40 and 50% for AdV41 (Fig. 5F/G). Infection of A549 cells appears to be clathrin- and actin -dependent despite an independence of EAdV overall uptake in presence of pitstop2 (Fig. 4A). This suggests additional clathrin-independent non-productive uptake pathways in A549 cells. In contrast, our uptake and infection assays for HuTu80 consistently suggest a role for CME in EAdV infection. Taken together, we conclude that EAdV can rapidly infect duodenal HuTu80 cells in a clathrin- and actin-dependent manner.

## Discussion

In this study, we aimed to evaluate the infectious pathway of EAdVs in duodenal HuTu80 compared to lung-derived A549 cells. Our results show that EAdVs have a kinetic advantage when infecting duodenal cells over lung-derived cells. Furthermore, we find that EAdV enter cells via a clathrin-dependent mechanism in A549 and HuTu80 cells, while clathrin-independent uptake was described for HEK293 cells. A faster and possibly more efficient uptake in HuTu80 cells is well in line with EAdV’s natural tropism in the gastrointestinal tract. Differential host cell requirements in uptake pathways and their translational implications have recently become obvious. For example, SARS-CoV-2 enters Vero cells by endocytosis and lung epithelial cells by fusion at the plasma membrane. Thus chloroquine has antiviral effects against SARS-CoV-2 in Vero cells, but not in cells of lung tissue origin (39). This underlines that infection models reflecting the target tissue are vitally important to understand mechanisms of virus entry and infection and to device strategies for antiviral treatment.

Historically, EAdV were fastidious and difficult to grow to high infectious titer in tissue cultures (8,9). Even to date, we observe that overall yields and infectious-to-non-infectious particle ratios are poorer than for other adenoviruses (e.g., AdV5 or AdV-D types) (unpublished results, Arnberg Lab, compare S1 Fig.). Despite their relevance as a leading cause of acute infant gastroenteritis (4), EAdV remain understudied (40). EAdVs are found in stool samples from children with acute gastroenteritis but we lack knowledge on their initial target cells or target regions in the gastrointestinal tract. Hence, virus entry studies originate from easily accessible cell lines like A549 and HEK293 cells. In these studies, EAdV uptake into A549 cells was slow, and a major fraction of particles were still neutralized by antibodies after 4 h (20). Similarly, AdV41 entry into HEK293 was inefficient (21). These important observations prompted us to study EAdV host factors in cells originating from the gastrointestinal tract. Our results show a faster uptake of EAdVs into HuTu80 cells when directly compared to A549 cells. As the gastrointestinal tract is specialized on nutrient resorption, it is tempting to speculate that the kinetic advantage lies in a different overall endocytic activity in HuTu80 cells. It is, therefore, critical to discriminate between productive and unproductive endocytic uptake of virus particles. To this end we not only compared global virus uptake but also uptake resulting in successful delivery of the virus DNA to the nucleus (i.e., infection). Consistently, we found a faster productive uptake of EAdVs in HuTu80 cells as compared to A549 cells, but no enhanced uptake kinetics for AdV5. This suggests that the infectious route of EAdVs rather than overall endocytic activity was enhanced in HuTu80 cells.

We hypothesized that the observed difference in internalization behavior between A549 and HuTu80 cells may result from cell-type-dependent surface interactions and/or differential availability of surface receptors, which could in turn affect particle mobility at the cell surface. For this reason, we tracked particles on the cell surface and analyzed their trajectories. We performed single particle tracking of EAdVs and AdV5 on HuTu80 and A549 at early time points after binding, when we suspected most particles to be on the cell surface based on our uptake and infectious internalization kinetics (Fig. 1). We found high variability in the measured diffusion coefficient for all conditions and motion types. We speculate that the variability arises from the multiple possible interactions that occur between AdVs and the cell surface. Strong multivalent interactions between cellular receptors and particle fiber proteins may cause immobile or slow diffusing confined particles, as it would be the case with particles in endocytic pits but also during interaction with attachment factors like heparan sulfate proteoglycans (HSPGs) after initial binding. Higher mobility of particles can arise from binding to mobile receptors on the cell membrane, either displaying free or confined diffusion, or being coupled with the underlying moving matrix. The latter coupling results in directed motion, as suggested for AdV2 in human embryonic retinoblast (HER) 911 cells (16). Matrix coupled motion was observed to be actin-dependent for other viruses (41–43) or even in a microtubule-assisted manner the C-type lectin DC-SIGN in dendritic cells (44). In addition, our method, although confining the observation to few hundreds of nanometers across the apical membrane of the cell, does not allow to distinguish between external and internalized particles. Thus, a small part of the observed motion is likely to be attributed to the movement of endosomes containing viral particles in the proximity of the membrane. We analysed the overall motion behaviour of each particle by developing the “average motion coefficient” (AMC). Here, we found that a higher fraction of the particle population showed a higher mobility with regards to surface explored (AMC score of 2). It is tempting to speculate that the greater fraction of mobile particles reflects viruses bound to mobile receptor species possibly scouting the surface for their site of internalization comparable to faster diffusion of AdV2 interacting with its receptor CAR prior to endocytic uptake (16). EAdV interact with their long fiber with CAR in Chinese hamster ovary cells expressing human CAR (14) and show affinity to HSPGs likely through their short fiber protein as confirmed in surface plasmon resonance measurements with purified fiber knobs (13). In addition, AdV41 penton base can engage laminin-binding integrins as co-receptors in human colon cancer cells (17), however, cell entry competent receptor interactions on A549 or intestinal cells remain unknown. In light of this and taking into account that HuTu80 cells carry less HSPGs but slightly more CAR on their cell surface than A549 cells (data not shown), EAdV may be slowed by multivalent interactions with HSPGs on A549 and have a surface sampling advantage leading to faster internalization in HuTu80 cells. This hypothesis is also supported by the finding that large extracellular domains, like HSPGs, have smaller diffusion coefficients on giant luminal vesicles (45). To validate this hypothesis, virus particles with short or long fibers only, would be valuable tools to assess the contribution of known receptors to the entry kinetics and diffusion behaviour. Studying surface interactions of EAdV on the two cell lines may also reveal cell type or tissue specific receptors that explain differences in the kinetic behaviours.

EAdVs enter the intestine by passing through the stomach with a low pH milieu. Previous studies demonstrated that EAdV particles are resistant to low pH explaining in part their tropism (18). In our study, we established a low pH wash protocol to block EAdV entry into intestinal epithelial cells. While this seems counterintuitive, we could reproducibly show that cell-bound EAdV particles become non-infectious upon low pH wash. We conclude that EAdV-cell surface molecule interactions induce a conformational change in the EAdV capsid, that renders the virus particles acid sensitive. This is in line with a sequence of entry cues for EAdV as seen for AdV2 in a step-wise process (16,46), starting with attachment to host factors followed by secondary low pH priming in endosomes, which then triggers endosomal membrane rupture and escape. Future work will address whether such sequential entry cues govern EAdV entry into the intestinal epithelium.

Here, we report the first TEM imaging of EAdV particles entering cells of physiological tissue origin. In TEM, EAdV increasingly localized to intracellular compartments (endosomal structures and cytosol) in both cell lines over time reflecting successful virus entry and endosomal escape. The effect was again more pronounced in HuTu80 cells and strongest after 2 h of internalization. This was in line with the effects seen in the early phases (0 – 6 hpi) in the infectious internalization kinetics and supporting the kinetic advantage observed. Virus particles were mainly observed on smooth membranes and membrane invaginations lacking a visible clathrin coat structure indicating a non-clathrin dependent entry pathway. We demonstrated, however, that productive virus uptake and infection was clathrin-dependent in inhibition and RNAi assays. This can be explained by EAdV’s slow and protracted entry, where single virus entry events likely occur asynchronously and over an extended period of time as seen for other viruses e.g. Human Papillomavirus 16 (47). In contrast, CME is a fast process, which occurs within 2 min (48). In our assays, we are therefore observing single internalization events at any time point rather than a synchronous entry of many particles. Those events themselves may be fast as they occur by CME. In static experiments like TEM, where we are observing cells at a fixed time point, we hypothesize that we likely miss most endocytic events. We observed a small fraction of viruses in endocytic pits mostly without a clathrin coat, which could indicate that those endocytic events are mechanistically slower or possibly delayed due to inefficient cargo recognition. Orthogonal methods including pharmacological perturbation and RNA interference are clearly better suited to reveal involvement of fast uptake mechanisms. Nonetheless, our TEM analysis confirms a faster uptake of EAdVs into cells of intestinal origin and our data collectively argues for including intestinal epithelial cell lines in the toolbox of EAdV studies.

Our pharmacological perturbation and RNA interference studies reveal clathrin-mediated endocytosis as the major entry pathway of EAdVs into intestinal cells. This is in contrast to previous studies on HEK cells (21) and to our observations in A549 cells, highlighting the importance of choice of experimental system. The effects during infectious internalization experiments with pitstop2 only showed a partial effect at the highest inhibitor concentration. Due to high cytotoxicity and possible off-target effects, the drug was washed out after half of the incubation time. This may be the reason for a lack of dose-dependency for pitstop2, as successful entry and hexon production may be possible after wash-out of the drug. Because of these limitations and the debatable specificity of pitstop2 (49,50), we additionally confirmed importance of CME for EAdV entry by RNA interference with the clathrin adapter AP2. Taken together, through orthogonal assays we demonstrated an important role of CME in productive EAdV entry into intestinal epithelial cells. As the cell system used is derived from tissue of EAdV tropism, we propose that CME is a physiological uptake route. Studies using human intestinal organoids need to confirm this notion.

Our study highlights the importance of studying host factors and infection pathways in appropriate model systems. In contrast to previously used respiratory epithelial cell models, EAdV infection of duodenal HuTu80 cells revealed that EAdV entry is clathrin-dependent and that enhanced infection kinetics may be a measure for infection efficacy. Clearly, cell lines from different tissue backgrounds will remain an important tool to identify and validate host factors during virus infections. This study characterizes duodenal cells as a novel tool for the study of EAdV and reveals previously neglected aspects of the cell entry mechanisms of these medically relevant viruses.

## Materials and Methods

### Cells, media, chemicals

A549 and HuTu80 cells cultured in DMEM (Sigma, #D5648) supplemented with 1x pen-strep (Gibco, #15140122) and 20 mM HEPES (Thermo Fisher Scientific, #BP310-500) (DPH) and 10% FBS (Hyclone, #SV30160.03) at 37°C and 5% CO_2_. For infections medium without FBS was used for incubation (DPH) and changed to medium with low FBS until fixation (DPH + 2% FBS). Small molecule inhibitors were pitstop2 (Abcam, #ab120687), dynasore (Abcam, #ab120192), cytochalasinD (Abcam, #ab143484), methyl-β-cyclodextrin (Sigma, C4555-1G). CsCl was from Sigma (C4036).

### Viruses

AdV-F40 (strain Hovix), AdV-F41 (strain Tak), and AdV-C5 (strain adenoid 75) were produced in A549 cells and purified as described previously (51). In brief, detached cells were pelleted at 800 rpm, supernatants were discarded, and cell pellets were suspended in DMEM. For cell lysis and virus extraction, cell suspensions were supplemented with an equal volume of Vertrel XF (Sigma; #94884) and shaken thoroughly. Phases were separated by centrifugation for 10 min at 3000 rpm. The upper aqueous phase was loaded onto a CsCl step gradient with the densities 1.27, 1.32, and 1.37 (bottom to top) for ultracentrifugation in a SW40 rotor (Beckman Coulter) for 2.5 h at 4°C and 25000 rpm. The virus containing band was extracted from the gradient and desalted on a NAP column (VWR, Cytiva, Illustra NAP) by elution in PBS, pH 7.4 and frozen at -80°C after addition of 10% glycerol.

### AF488-labeled AdV

AdV stocks were produced as above and used frozen (AdV5) or fresh (AdV40 and AdV41) for covalent labeling with AF488-NHS (Thermo Fisher Scientific; #A20000). Viruses were used at a concentration of at least 500 ng/µL (best 1000 ng/µL) and incubated with 10-fold molar excess dye over virus in a 500 µL reaction for 1 h rotating and light protected at room temperature. Virus was separated from free dye by ultracentrifugation as above on a SW60Ti rotor (Beckman Coulter). The virus band was extracted and desalted on a NAP column (VWR, Cytiva, Illustra NAP) by elution in PBS, pH 7.4 and frozen in small aliquots at -80°C after addition of 4% glycerol.

### Immunofluorescence staining

75000 A549 cells were seeded in growth medium on ibidi 8-well µ-slides (ibidi, #80826) 24 h prior to experimentation. AF488-labeled AdVs were bound to cells at 4°C on ice for 1 h. Medium was exchanged, and cells were either kept at 4°C or shifted to 37°C for 4 h for internalization. Samples were transferred to 4°C for staining with anti-AF488-antibody (rabbit, Thermo Fisher Scientific; #A-11094) for 30 min on ice shaking, washed once with cold PBS + 2% FBS and subsequently stained with anti-rabbit IgG-AF568 for 30 min at 4°C shaking. Cells were fixed with 4% PFA/PBS for 20 min and stained with AF647-Wheat Germ Agglutinin (WGA, Thermo Fisher Scientific, # W32466) and Hoechst33342 (Thermo Fisher Scientific; #62249). Samples were imaged as z-stacks with a 63x objective on a Leica SP8 confocal microscope at the Biochemical Imaging Center Umeå (BICU). Images were processed by maximum intensity projection with Fiji (52).

### Uptake assay

Cells were detached with PBS/EDTA, pelleted, suspended in DPH + 10% FBS and kept in suspension on a shaker for 1 h for recovery. 2×10^5^ cells/well were pelleted in a v-bottom 96-well plate and washed with cold DPH. For uptake assays in presence of inhibitors, cells were preincubated with inhibitor dilutions in DPH for 30 min before subsequent virus incubation. Cells were resuspended in 100 µL DPH (with or without inhibitors) containing AF488-labeled viruses and incubated on ice for 1 h. Cells were pelleted and washed once with 100 µL cold DPH. Control samples were left on ice and internalization samples were transferred to 37°C for 4 h. Cells were pelleted and incubated with anti-AF488-antibody (rabbit, Thermo Fisher Scientific; #A-11094) for 30 min on ice shaking, washed once with cold PBS + 2% FBS and subsequently incubated with anti-rabbit IgG-AF568 for 30 min at 4°C shaking. Cells were washed once with cold PBS + 2% FBS and resuspended in 50 µL cold PBS + 2% FBS. Samples were measured by flow cytometry (BD Accuri) to assess mean fluorescence intensities of AF488 and AF568.

### Infectious entry assay

25000 A549 or 30000 HuTu80 cells per well were seeded in a black optical bottom 96-well plate (Greiner Bio-one; #655096) 24 h prior to experimentation. *Titration:* Virus stocks were titrated for every batch to determine a dilution that results in 20% absolute infection. Virus stocks were diluted serially in DPH, 50 µL of the virus dilution was added to cells and incubated for 2 h at 37°C. Inoculum was removed, samples were supplemented with DPH + 2% FBS, and incubated until 48 hpi. Cells were fixed by addition of 8% paraformaldehyde in PBS to the culture medium and incubated at room temperature for 15 min. *pH-wash:* pH sensitivity of EAdV infection after cell surface binding was tested by a 2 min pH wash after virus binding in DPH for 1 h on ice. Cells were incubated for 2 min with either 0.1 M citrate/PBS buffer (pH 3-6) or 0.1 M CAPS/PBS (pH 8-11) or PBS (pH 7.4). Samples were neutralized by two washes with PBS and subsequent addition of DPH + 2% FBS until fixation with 8% paraformaldehyde in PBS at 48 h post infection. *Kinetics:* Cells were washed once with DPH without FBS and transferred to 4°C. Medium was removed and 50 µL virus dilution was added to each well for binding at 4°C for 2 h. Inoculum was removed and warm DPH + 2% FBS was added to each well. At each time point, duplicate samples were washed with 0.1 M citrate buffer pH 3 for 2 min, subsequently washed gently with PBS and supplemented with warm DPH + 2% FBS until fixation with 4% paraformaldehyde in PBS at 48 h post infection. *Inhibitor:* Cells were incubated with inhibitors dilutions in DPH for 30 min at 37°C. Meanwhile, virus dilutions were mixed with the inhibitors dilutions in a dilution plate. Medium was removed from the cells and 50 µL virus-inhibitor dilution was added to each well for initial binding and internalization for 2 h at 37°C. Then, inoculum was removed and 100 µL inhibitor dilutions mixed with DPH + 4% FBS was added to each well for 24 h. Thereafter, medium was removed and 100 µL warm DPH + 2% FBS was added until fixation with 4% paraformaldehyde in PBS at 48 h post infection*. Staining:* Cells were permeabilized with 100% ice-cold methanol for 20 min at -20°C and rehydrated by washing with PBS and then stained for viral hexon protein expression with an anti-hexon primary antibody (mouse; Merck; #MAB8052) and an anti-mouse secondary antibody coupled to AF488 or AF647 (Thermo Fisher Scientific; #A-11001/#A-21235). Nuclei were stained with Hoechst33342 (Thermo Fisher Scientific; #62249) and used to determine cell numbers. Plates were scanned in a Biotek Cytation5 automated microscope with a 10x objective with subsequent image analysis for cell count and infection counting or using the Tina program a TROPHOS plate reader (Luminy Biotech Enterprises, Marseille, France).

### SiRNA knockdown

250000 A549 or 350000 HuTu80 cells per well were seeded in 12-well plates 24 h prior to experimentation. Cells were transfected with 15 nM siRNA (AP2M1: Thermo Fisher Scientific # 1299001, HSS101955; SCR: Thermo Fisher Scientific #12935300) with 3.33 µL Lipofectamine RNAiMax (Thermo Fisher Scientific, #13778075) following the recommended protocol. The transfection mixture was added dropwise to wells containing 1 mL DPH + 10% FBS and not removed until next day. Cells were split into a 6-well plate on the next day. Cells were transfected with 5 nM siRNA (A549) or 15 nM siRNA (HuTu80) with 5 µL Lipofectamine RNAiMax following the recommended protocol. The transfection mixture was added dropwise to wells containing 2 mL DPH + 10% FBS and kept until next day. Cells were seeded into 96 well plates for infection experiment as described above and into 48 or 24 well plates for Western Blot. Cells were harvested for Western Blotting by addition of 2x loading dye/DTT. Samples were boiled for 10 min and frozen at -20°C. Samples were separated on precast gel (4-12% NuPAGE Bis-Tris, Thermo Fisher Scientific, #NP0321BOX) with 1x MOPS buffer (Novex, NuPage, #NP0001) for 1 h at 100 V. Proteins were transferred to a nitrocellulose membrane (0.2 µm, BioRad, #162-0112) with transfer buffer (20% methanol, 39 mM glycine, 48 mM Tris base, 0.037% SDS) for 75 min at 100 V. Membranes were blocked with 2% ECL prime blocking reagent in PBS-T (Amersham; #RPN418) and incubated against AP-2 (mouse-anti-AP50, BD, #A611351) and GAPDH (rabbit-anti-GAPDH, Sigma, #G9545-100uL) in blocking buffer over night at 4°C. Signals were detected by chemiluminescence using anti-mouse/rabbit-HRP secondary antibodies (Novex, A16072/A16104) and ECL substrates (Thermo Fisher Scientific, #34577/34095). Membranes were imaged on an Amersham imager 680 (GE/Cytiva).

### Transmission electron microscopy

1×10^6^ A549 and HuTu80 cells per well were seeded in a 6-well plate on the day prior to experimentation. Cells were transferred to ice and washed twice with cold DPH. Viruses were diluted at 50000 particles per cell (based on 11 ng virus preparation contains 4×10^7^ particles (53)) and added to cells on ice. Samples were incubated on ice for binding for 1 h shaking. Thereafter, virus inoculum was removed and warm DPH + 2% FBS was added to each well. Samples were transferred to 37°C and internalization was allowed for 12.5 min, 2 h and 6 h. After incubation, medium was removed and 1.5 mL EM-fixative (2.5% glutaraldehyde, 0.05% malachite green, 0.1 M phosphate buffer (PB; NaH_2_PO_4_ + MQ-water) (Taab Laboratory Equipment Ltd, Aldermaston, UK). for 1 h at room temperature. Fixed samples were washed twice with 2 mL 0.1 M PB and kept in 0.1 M PB until processing. All the below chemical fixation steps were performed in the microwave Pelco biowave pro+ (Ted Pella, Redding, CA) unless stated. The samples were rinsed 2x with PB and post-fixed with 0.8% K_3_Fe(CN)_6_ and 1% OsO_4_ in 0.1 M PB for 14 min and were rinsed 2x with MQ-water. The samples were then stained with 1% aqueous tannic acid. Following 2x rinse in MQ-water, samples were stained with 1% aqueous uranyl acetate (Polysciences, Inc., Hirschberg an der Bergstrasse, DE). After 2x MQ-water rinses, samples were dehydrated in gradients of ethanol (30%, 50%, 70%, 80%, 90%, 95%, 100% and 100%). The samples were infiltrated with graded series of Spurr resin in ethanol (1:3, 1:1 and 3:1). After microwave, samples were infiltrated twice with 100% resin before polymerization overnight at 60°C. Samples were sectioned (70 nm) and imaged with Talos 120C (FEI, Eindhoven, The Netherlands) operating at 120 kV at the Umeå Core Facility for Electron Microscopy (UCEM). Micrographs were acquired with a Ceta 16M CCD camera (FEI, Eindhoven, The Netherlands).

### Diffusion measurement

Imaging of AdV particles diffusing on live cells was performed using a Nikon Ti2-E inverted microscope equipped with a two-color laser source (wavelengths: 488 nm and 562 nm), a Prime 95B sCMOS camera (Teledyne Photometrics) and a multiband filter cube (86012v2 DAPI/FITC/TxRed/Cy5, Nikon Corp.). The images were collected using a 60x oil immersion objective (numerical aperture: 1.49, MRD01691, Nikon Corp.), 488 nm excitation wavelength and HILO microscopy configuration, which allows imaging of the apical cell surface while reducing out-of-focus signal and background noise when compared to epifluorescence. For the experiments, 40000 A549 and 50000 HuTu80 cells per well were seeded in a glass-bottom 96-well plate 24 h before imaging. The following day, the culture medium was removed and replaced with warm DMEM without phenol red and 2% FBS to reduce the background signal in fluorescence imaging. Cells were incubated at 37°C for 1 h and, right before the experiment, the plate was transferred under the microscope and kept at 37°C and 100% humidity in a dedicated on-stage incubator (Okolab). 100 µl of AF488 labelled AdV in a premixed 1:50 dilution was added in each well, pipetting thoroughly to homogenize the virus particle distribution in the cell medium. After 5 min, the cell supernatant was removed, and the well was washed twice with fresh prewarmed medium. Cells were imaged 5 and 30 minutes after washing. Videos were acquired at 10 frames per second for 2 min. Two videos were acquired for the 5 min and between 2 and 4 videos for the 30 min time point. The frame rate was optimized to guarantee good time resolution and a high signal-to-noise ratio. All conditions were acquired in duplicates.

### Single particle tracking and diffusion analysis

The recorded time-lapses were processed to remove the background and improve contrast with an in-house ImageJ script. First, a Gaussian filter with a sigma of 0.5 pixels was convolved to the image to reduce the camera noise. Bleaching was then corrected using a built-in Fiji function, which fits the variation of total intensity in time to an exponential fit (52). The background signal was calculated by averaging over all the frames and then applying a Gaussian filter with a sigma of 20 pixels. This removes the fine details, like immobile viruses, while maintaining the broader cellular features. The background was then subtracted from each frame. The virus particles were then detected and tracked using the Laplacian Gaussian filter in the TrackMate plugin, which allows for sub-pixel resolution on the particle position (54). Linking between frames is performed using a maximum linking displacement of 1 µm, and a maximum frame gap of 10. The obtained trajectories were segmented according to the characteristics of the motion displayed using a DC-MSS algorithm (24), and each segment was assigned to a motion type, according to the slope of the moment scaling spectrum. The motion type considered were immobile, confined motion, free diffusion and directed motion. In-house scripts were used to extract diffusion parameters, e.g. Diffusion coefficient (D) and confinement radius (R_C_), of both each segment and each unsegmented track. Only tracks with a duration of more than 300 frames (30 seconds) were considered in the analysis of unsegmented tracks. The tracks were classified considering the average time spent in each motion type. A numerical value was assigned to each motion type (0: immobile, 1: confined motion, 2: free diffusion, 3: directed motion) and the weighted average over the time spent in each motion type was calculated resulting in a numerical coefficient (“average motion coefficient”) between 0 (fully immobile) and 3 (only directed motion). A value of 1.5 was considered as the threshold between particles dominated by confined diffusion and particles displaying mostly free or directed motion.

## Acknowledgements

We acknowledge the Biochemical Imaging Center and Umeå Core Facility for Electron Microscopy at Umeå University and the National Microscopy Infrastructure, NMI (VR-RFI 2019-00217) for providing assistance in microscopy.

## Supporting information

**S1 Fig. Infection titration A549 and HuTu80.** A-C) Susceptibility of A549 (black) and HuTu80 (red) cells to AdV40, AdV41, and AdV5 infection. Shown are typical dilution series and absolute infection values from three independent experiments. Error bars indicate the standard error of the mean.

**S2 Fig. Low pH wash after cell binding renders EAdVs non-infectious.** A/B) Sensitivity of EAdV infection to low pH wash after binding to A549 cells. Error bars indicate the standard error of the mean. Statistical significance determined using two-way ANOVA test: *: p<0.05, **: p<0.01, ***: p<0.001, ****: p<0.0001.

**S3 Fig. No clear trend is observed in the diffusion coefficient.** Diffusion coefficient of AdVs after 5 and 30 min from the removal of the inoculum and on A549 and HuTu80 cells for (A) confined diffusion, (B) free diffusion and (C) directed motion. Grey dots indicate the mean value for each recorded video. Error bars indicate the standard error of the mean.

**S4 Fig. Diffusion regimes segmentation:** A) Fraction of the total track time spent in each motion type by AdV particles on A549 and HuTu80 cells after 5 and 30 min from the removal of the inoculum. B/C) Comparison of the total track time spent in immobile (B) and directed (C) diffusion on A549 and HuTu80 cells after 5 and 30 min. Grey dots indicate the value for each recorded video. Error bars indicate the standard error of the mean. Statistical significance determined using two-way ANOVA test: *: p<0.05, **: p<0.01, ***: p<0.001, ****: p<0.0001. D) Example track showing how the maximum distance travelled per track segment was calculated. E) Comparison between the fraction of time, of the total length of the track in each segment, and of the maximum distance traveled, i.e., the distance between the two furthest points of each segment, in each motion regime for the track in (D). F) Fraction of the distance travelled in each motion regime for AdV particles on A549 and HuTu80 cells.

